# Clonally Expanded CD8 T Cells in AML Adopt an NK-Like Program Distinct from Canonical Exhaustion

**DOI:** 10.64898/2026.09.30.755147

**Authors:** Yunli E Chu, Matthew A Cottam, Chad R Potts, Thomas Gracie, Ciarra Polsley, Justin A Cartailler, Michael R Savona, Rui Lu, Stanley C Lee, Robert Welner, P Brent Ferrell

## Abstract

CD8 T cells are key mediators of anti-tumor immunity, yet their differentiation states and clonal architecture within the acute myeloid leukemia (AML) bone marrow microenvironment remain incompletely defined. We performed high dimensional immune profiling of bone marrow samples from 46 newly diagnosed AML patients using mass cytometry to probe the microenvironment and T cell states. We used single-cell RNA and T cell receptor (TCR) sequencing to define CD8 T cell phenotypes, transcriptional states, and clonal architecture. AML-associated CD8 T cells were skewed toward terminally differentiated effector memory cells re-expressing CD45RA (TEMRAs), while canonically exhausted T cell populations were rare. Functional studies revealed that TEMRA CD8 T cells robustly produced IFNγ and TNFα upon in vitro stimulation. Clonal expansion was concentrated within the TEMRA compartment. Comparison of TCR sequences with known viral epitopes identified a small subset of expanded viral-specific clonotypes, while the majority of highly expanded clonotypes had unknown specificity and may include AML-specific T cells. Differential gene expression analysis revealed shared cytotoxic effector programs during clonal expansion but diverging activation signatures: viral-specific clones upregulated canonical T cell receptor signaling genes, whereas expanded clones of unknown specificity exhibited enrichment of natural killer (NK) receptor-associated genes. These genes comprised the leukemia-associated NK-immune (LaNKi) score which was elevated in an independent AML dataset compared to pan cancer tumor infiltrating lymphocytes. These findings define the CD8 T cell differentiation landscape and clonal architecture in AML marrow at diagnosis and suggest that T cell dysfunction in AML diverges from classical activation and exhaustion paradigms.

**Key points:**

1. CD8 T cells in AML are enriched for terminally differentiated states not defined by traditional checkpoint-inhibitor expression.
2. The LaNKi score marks clonally expanded CD8 T cells of unknown specificity and is validated across independent AML and pan-cancer datasets.

## Introduction

Acute myeloid leukemia (AML) is the most common acute leukemia in adults and carries a poor prognosis, with a 5-year survival rate of 32.9%^1^. Despite advances in targeted therapy and stem cell transplantation, most patients will relapse and die^2–4^. Unfortunately, the success of immunotherapies in cancers such as melanoma, small cell lung cancer, and lymphoma^5–7^ have yet to translate to meaningful T cell-targeting approaches in AML^8^, highlighting the need for a deeper understanding of T cell biology in this disease.

CD8 T cells are powerful mediators of anti-tumor immunity^9^, yet few studies have systematically profiled CD8 T cells within the AML bone marrow microenvironment at the resolution needed to link surface phenotype, transcriptional state, and clonal identity. Prior reports demonstrate that CD8 T cells in AML are skewed toward a terminally differentiated, senescent-like state marked with NK-like features and clonal expansion. These studies also report a lower abundance of naïve CD8 T cells. In contrast to solid tumors, canonical exhaustion phenotypes defined by checkpoint inhibitor expression are rare in AML^10–14^. These features may contribute to the limited success of immunotherapy.

In solid tumors, CD8 T cells infiltrating the tumor mass, commonly called tumor-infiltrating lymphocytes (TILs), serve as a key biomarker of immune engagement^9,15^. In contrast, the BM is both the site of leukemic transformation and a natural reservoir for naïve and memory T cells under healthy, steady-state conditions^16,17^. Consequently, CD8 T cells detected in AML marrow may represent a mixture of leukemia-specific populations and bystander cells that were resident in or circulating through the marrow prior to disease onset, meaning expanded clones cannot be assumed to be leukemia-reactive by proximity alone. Disentangling these populations is critical to understanding anti-leukemic immunity and identifying targets for immunotherapy, yet no study has directly compared the transcriptional programs underlying the clonal expansion of viral-specific bystander clones versus clones of unknown specificity.

Previous studies of T cell responses in AML have been limited by narrow phenotypic panels, small patient cohorts, and a primary reliance on transcriptomics. To address these gaps, we profiled BM CD8 T cells from a large cohort of patients with newly diagnosed AML using orthogonal proteomic, transcriptomic, and TCR sequencing approaches. This multimodal dataset enables the integration of surface phenotype, transcriptional state, and clonal architecture at single-cell resolution. We further investigate how leukemia phenotype shapes the CD8 T cell compartment and compare viral-specific T cell responses to those with unknown specificity define the molecular programs associated with clonal expansion. Together, these analyses provide a comprehensive framework for understanding CD8 T cell states in AML.

### Methods (also see Supplemental Methods) Sample collection

Patients with newly diagnosed AML at Vanderbilt University Medical Center provided written informed consent prior to enrollment. BM collection and analyses were conducted under protocols approved by the Vanderbilt University Medical Center Institutional Review Board (IRB #171654) and followed the ethical standards of the institution and the guidelines of the Declaration of Helsinki. BM mononuclear cells (BMMCs) were isolated by density-gradient centrifugation, cryopreserved in fetal bovine serum supplemented with 10% dimethyl sulfoxide, and stored in liquid nitrogen.

### Mass cytometry

BMMCs were processed in batches of up to 20 samples to minimize batch effects. For the T cell-specific panel, T cells were first negatively enriched from BMMCs (STEMCELL Technologies). Cells were stained with cisplatin (Standard BioTools), followed by fixation in 0.4% paraformaldehyde. Samples were palladium-barcoded (Standard BioTools), and pooled before incubation with extracellular antibody cocktails^18^. For panels incorporating intracellular markers, cells were fixed and permeabilized with the Foxp3/Transcription Factor Staining Buffer kit (Tonbo Biosciences) and stained with intracellular antibodies. Mass cytometry antibodies are listed in Supplemental Table 1.

Following antibody labeling, cells were fixed in 1.6% paraformaldehyde and permeabilized with ice-cold methanol. Methanol-permeabilized samples were stored at -80°C. Prior to acquisition, samples were incubated with an iridium-based DNA intercalator (Standard BioTools), and resuspended in Cell Acquisition Solution (Standard BioTools) containing EQ Four Element Calibration Beads (Standard BioTools). Data were acquired on a Helios mass cytometer (Standard BioTools).

### Gene signature scoring

UCell-v2.14^32^ was applied using exhaustion genes (*CTLA4, HAVCR2, LAG3, PDCD1, TIGIT,* and *ENTPD1*), activation/proliferation genes (*NME1, IL2RA, CD38, HLA-DRA, HLA-DRB1, HLA-DRB5, MKI67,* and *PCNA*), and genes from the LaNKi signature (*HOPX, KLRC4, KLRK1, KLRG1, and NFKBIA*).

### Annotating TCR antigen-specificity

To identify putative viral-specific clones, we cross-referenced CDR3-TCRΒ sequences against the VDJdb curated database of published TCR sequences and epitopes^33^. Results were restricted to MHC class I-restricted epitopes. Matched clones were annotated as viral-specific.

### T Cell stimulation

BMMC samples were rested in RPMI-1640 (Gibco) supplemented with 10% fetal bovine serum (Cytiva), 1% penicillin-streptomycin (Gibco), 1% HEPES (Gibco), 1% glutamax (Gibco), and 50μM beta-mercaptoethanol (Gibco) at 37°C. Cells were seeded at 1M cells/ml and stimulated with vehicle (dimethyl sulfoxide, Gibco), Dynabeads Human T-Activator αCD3/αCD28 beads (Thermo Fisher Scientific), 20 ng/ml phorbol 12-myristate 13-acetate (PMA) and 1μg/ml ionomycin (Invitrogen). All conditions contained 1x brefeldin A (eBioscience) and 1x monensin (eBioscience) and were incubated at 37°C for 6 hours.

### Spectral flow cytometry

Samples were stained with the extracellular antibody cocktail (Supplemental Table 3), fixed and permeabilized using the Foxp3/Transcription Factor Staining Buffer kit, and stained with the intracellular antibody cocktail (Supplemental Table 3). Samples were stored at 4°C until acquisition on a Cytek Aurora flow cytometer.

### Statistical analysis

Nonparametric tests were used, including the Wilcoxon rank sum test and Kruskal-Wallis test. P values less than 0.05 were considered significant. Pearson correlation was used for correlation analyses. To evaluate the association between effector memory CD8 T cell frequency and patient survival, we used Cox proportional hazards regression with overall survival as the outcome and effector memory CD8 T cell percentage as a continuous predictor, adjusting for patient age and ELN risk status.

## Results

### Mature immune cells are rare in AML

We collected BM samples from newly diagnosed AML patients (N=46) spanning diverse mutational and cytogenetic backgrounds (Supplemental Figure 1A, Supplemental Table 4), as well as BM samples from healthy controls (n=5). Each sample was profiled using an AML-focused mass cytometry panel to characterize the whole BM compartment and a complementary T cell-specific panel to resolve T cell phenotypes. Based on progenitor and lineage marker expression, we identified nine major cell clusters in whole marrow (Supplemental Figure 1B-C). The broad “HSPC-myeloid” compartment encompassed four of these clusters: HSC-like, progenitor-like, monocyte/macrophage-like, and myelocyte-like populations.

As expected, AML samples showed a marked reduction in the abundance of mature immune lineages, including B cells, T cells, NK cells, and erythroid populations (Supplemental Figure 1D-E), reflecting the overrepresentation of the HSPC-myeloid compartment, which captures both leukemic blasts and healthy progenitors. This highlights the reduction of bystander immune populations in the leukemic niche.

When comparing lineage frequencies with clinical outcomes, only the frequency of BM CD8 T cells showed a significant association with remission status (Supplemental Figure 1F). Patient age did not correlate with CD4 or CD8 T cell abundance (Supplemental Figure 1G-H), nor with remission outcomes (Supplemental Figure 1I). Finally, neither CD4 nor CD8 T cell frequencies were associated with FLT3, DNMT3A, NPM1, CEBPA, TET2, or NRAS mutation status (Supplemental Figure 2A-B), suggesting that global T cell abundance is not shaped by specific mutational drivers. Though the reduction of T cells may be purely due to increased AML burden, the increased CD8 T cell abundance in remission cases suggests these immune cells may have an important role in treatment response.

### AML-associated CD8 T cells are enriched for TEMRAs

Given that CD8 T cells are the primary effectors in the immune response to cancer, we next sought to dissect the CD8 T cell compartment in greater detail. Using the T cell-specific mass cytometry panel, we subclustered BM CD8 T cells into six clusters based on canonical differentiation markers (Figure 1A-B). Compared to healthy controls, AML samples showed a marked reduction in naïve CD8 T cells. Central/resident memory, stem cell memory, and effector memory frequencies were largely unchanged. In contrast, both early and late TEMRA populations were significantly enriched in AML, indicating a skewing toward terminal differentiation (Figure 1C-D).

**Figure 1.**
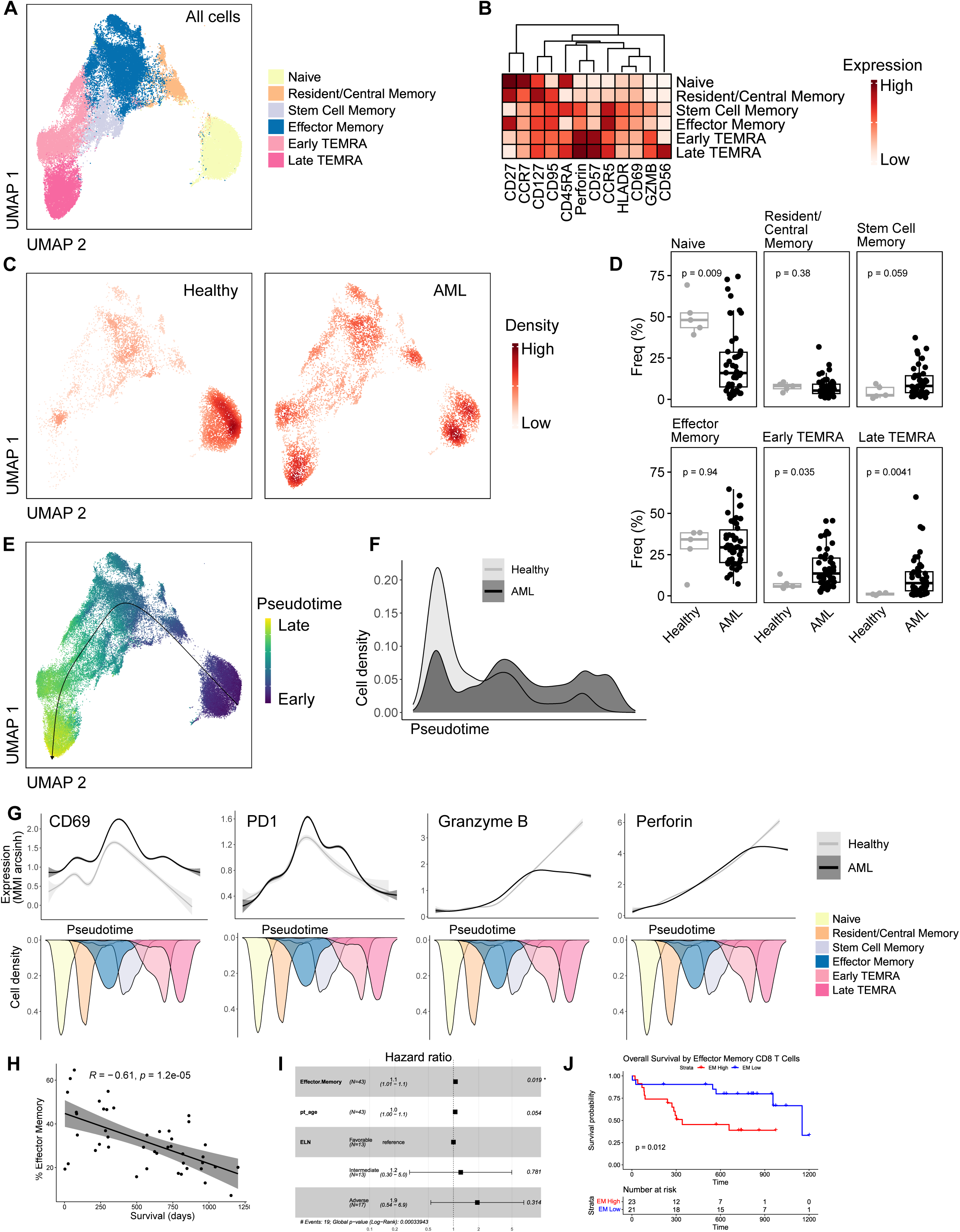
Dissecting the AML-associated CD8 T cell landscape. A) UMAP of CD8 T cells from all samples colored by cell type (N=51, #cells=66,971). B) Protein markers used to define cell types. C) UMAP of CD8 T cells from healthy (left, N=5, #cells=8,162) and all AML (right, N=46, #cells=8,162 randomly down sampled) marrows colored by density. D) Frequency of Naive, Resident/Central Memory, Stem Cell Memory, Effector Memory, Early TEMRA, and Late TEMRA CD8 T cells in healthy and AML marrows. Frequencies are calculated as a percentage of CD8 T cells. P-values calculated using Wilcoxon test. E) UMAP of CD8 T cells from all samples colored by calculated pseudotime with trajectory arrow spanning from early to late pseudotime. Pseudotime calculated using the Slingshot package with naïve cluster as the root node. F) Cell density plot of CD8 T cells from healthy (light grey) and AML (dark grey) marrows across pseudotime. G) Protein expression of CD69, PD1, granzyme B, and perforin across pseudotime of healthy (light grey) and AML (dark grey) associated CD8 T cells. MMI arcsinh = arcsinh transformed median mass intensity measured by mass cytometry. Mirrored density plots depict the dominant cell types across pseudotime. H) Pearson correlation between percentage of effector memory CD8 T cells and patient survival. I) Multivariable Cox proportional hazards model for overall survival in the AML cohort. J) Kaplan-Meier plot of overall survival of AML patients dichotomized by effector memory frequency above vs below cohort median. P-value calculated using log-rank test.

To evaluate differences across the continuum of differentiation states, we performed pseudotime analysis. The trajectory recapitulated the expected naïve to TEMRA progression (Figure 1E), with AML samples displaying a pronounced density shift toward late-differentiated states (Figure 1F). Along this continuum, several activation and effector genes differed between healthy and AML-associated CD8 T cells. CD69 expression was uniformly higher in AML CD8 T cells across pseudotime, whereas PD-1 levels were comparable overall but modestly elevated within effector memory and early TEMRA states. Granzyme B and perforin increased with differentiation in both groups, yet AML CD8 T cells demonstrated attenuated cytotoxic molecule expression at the terminal end of the trajectory (Figure 1G).

Effector memory T cell abundance was anti-correlated (R=-0.61, p=1.2e^-05^, Figure 1H) with survival. Effector memory frequency increased with age and naïve frequency decreased with age, while other cell types were not significantly correlated with age (Supplemental Figure 3A-F). However, in a multivariable Cox proportional hazards model for overall survival, adjusting for patient age and ELN risk status, effector memory T cell abundance was independently associated with survival outcome (HR=1.10, p=0.019; Figure 1I). Kaplan-Meier analysis demonstrated that patients with high effector memory abundance (above cohort median) had significantly lower overall survival compared to those with low effector memory abundance (below cohort median) (log rank p=0.012; Figure 1J). Taken together, these findings suggest that effector memory frequency in the CD8 T cell compartment is negatively associated with overall survival.

### AML-associated TEMRAs have preserved cytokine production capabilities

CD8 TEMRA cells have been previously described in the contexts of chronic inflammation including cancer. In these contexts, they are characterized by terminal differentiation and expression of cytotoxic programs despite loss of costimulatory receptors^34–39^. In AML BM, CD8 early and late TEMRAs expressed high levels of CD57, a marker associated with senescence^40^ (Supplemental Figure 4A). Prior research indicates that senescent CD8 T cells are not a defective end-stage population. Instead, they may rely on TCR-independent mechanisms for activation^41^. Based on this phenotype, we sought to assess the cytokine-producing capacity of AML-associated TEMRAs.

To evaluate effector function, a subset of BM samples from AML patients and healthy donors were stimulated *in vitro* with vehicle, αCD3/αCD28 beads, and PMA and ionomycin. Samples were analyzed by flow cytometry (gating scheme in Supplemental Figure 4B) for intracellular IFNγ and TNFα and degranulation marker CD107a. CD8 TEMRAs from AML patients robustly produced IFNγ and TNFα, with frequencies comparable to those observed in healthy donor TEMRAs (Supplemental Figure 4C-D). Degranulation potential was also comparable in TEMRA populations (Supplemental Figure 4E). These findings of preserved cytokine production and degranulation capabilities combined with decreased cytotoxic molecule expression shown in pseudotime analysis suggest an imbalance in CD8 T cell effector functions in AML.

### Select AML subtypes are associated with T cell landscape

Given the heterogeneity of AML and the possibility that leukemic biology may shape the immune microenvironment, we next asked whether specific CD8 T cell phenotypes were associated with particular AML subtypes. Because leukemic blasts are contained within the broad “HSPC-myeloid” cluster, we further resolved this compartment into eight phenotypically distinct subsets: HSC-like, CD56⁺ HSC-like, CD56⁺ early progenitor-like, myeloid progenitor-like, late myeloid progenitor-like, CD36⁺ monocyte-macrophage-like, CD4⁺ monocyte-like, and myelocyte-like populations (Figure 2A-B). The relative abundance of these subsets varied widely across patients, underscoring the phenotypic diversity of AML (Figure 2C).

**Figure 2.**
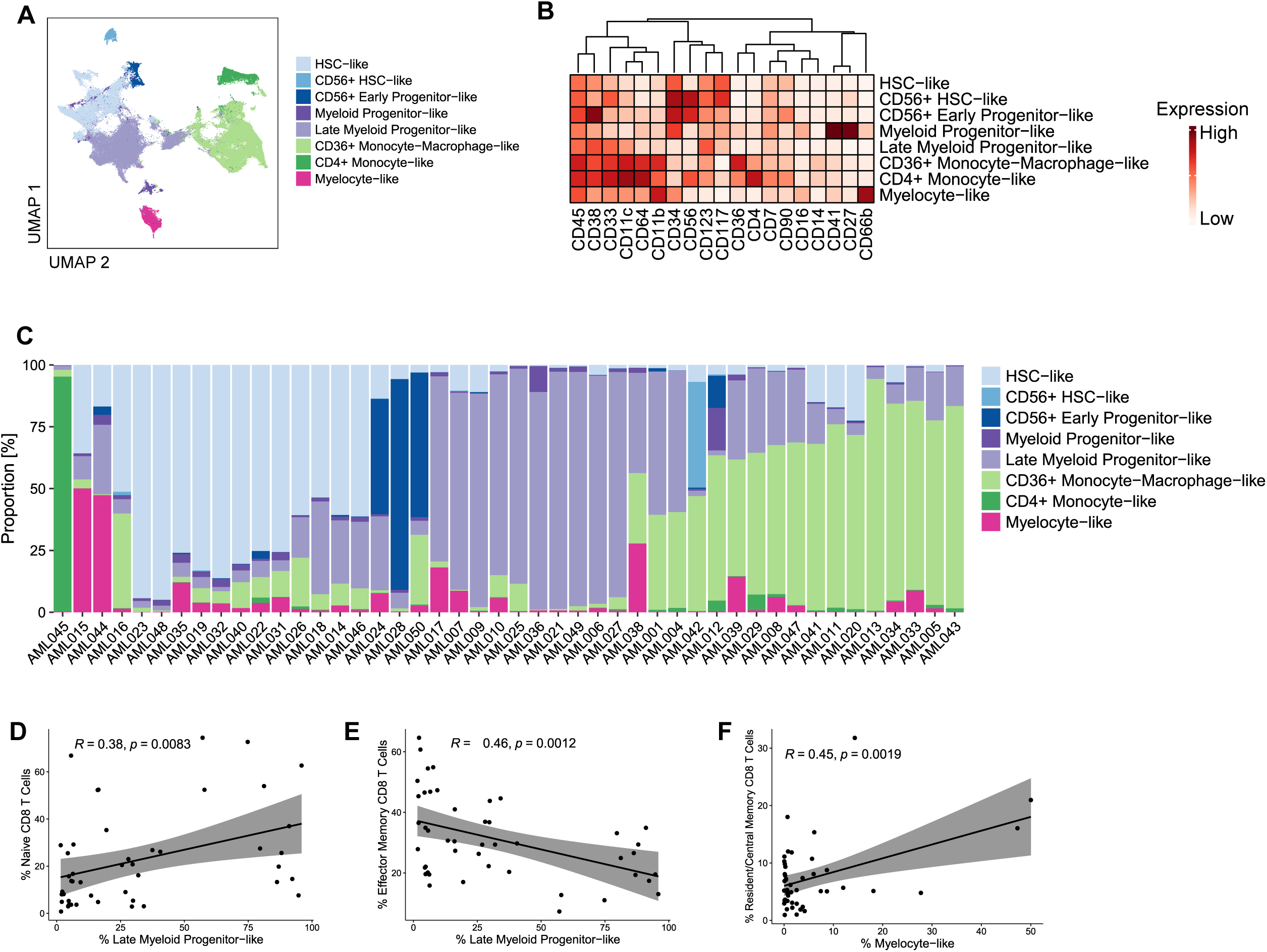
AML phenotype correlates with CD8 T cell phenotype. A) UMAP of HSPC-myeloid cells from AML marrows colored by cell type (N=46, #cells=412,697). B) Protein markers used to define cell types. C) Proportion of each cell type across AML samples. D) Pearson correlation between percentage of Naive CD8 T cells and percentage of Late Myeloid Progenitor-like cells. E) Pearson correlation between percentage of Effector Memory CD8 T cells and percentage of Late Myeloid Progenitor-like cells. F) Pearson correlation between percentage of Resident/Central Memory CD8 T cells and percentage of Myelocyte-like cells.

We next examined how these AML phenotypic states related to variation in the CD8 T cell landscape. The late myeloid progenitor-like population was positively correlated with higher naïve CD8 T cell frequencies but inversely correlated with effector-memory CD8 T cells (Figure 2D-E). In addition, the myelocyte-like subset was positively correlated with resident/central memory CD8 T cells (Figure 2F). Together, these findings indicate that late myeloid progenitor-like cells may be associated with impaired T cell priming or activation, resulting in preservation of the naïve T cell pool and reduced differentiation into effector memory cells. Myelocyte-like cells may support the retention of resident/central memory CD8 T cells natively found in the BM under steady-state conditions. These observations raise the possibility that distinct AML phenotypes can shape the T cell landscape through differential antigen presentation and priming, cytokine production, or cell type specific interactions.

### Canonically exhausted T cells are rare in AML

To probe transcriptional and clonal properties of CD8 T cell states in AML, we performed single-cell RNA and TCR sequencing on a subset of 16 patient samples. Based on transcriptomic features, CD8 T cells (n=23,445 cells) segregated into eight clusters: naïve, mucosal-associated invariant T cells (MAIT), activated, exhausted-like, central memory, effector memory, GZMK TEMRA, and GZMB TEMRA populations (Figure 3A-B). Antibody-derived tags for CD45RA and CD45RO aided in the identification of TEMRA populations. Using canonical activation and exhaustion gene signatures^42^, we found that activated/proliferating and exhausted-like phenotypes were present but comprised only small fractions of the CD8 T cell compartment (0.98% and 1.24%, respectively, Figure 3C-D).

**Figure 3.**
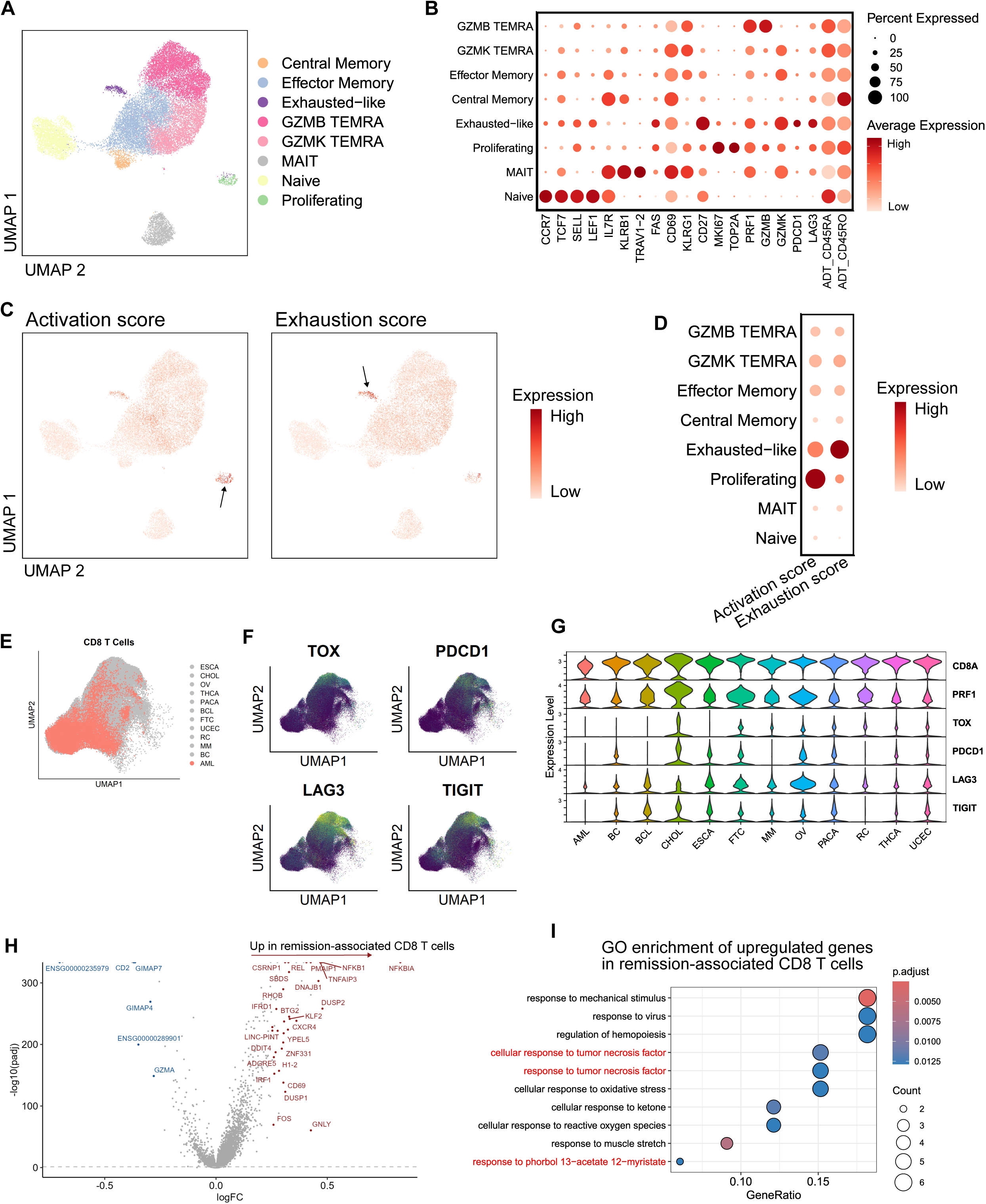
Single cell sequencing reveals a remission-associated T cell activation signature. A) UMAP of CD8 T cells from single cell RNA sequencing of AML marrows, colored by cell type (N=16, #cells=23,445). B) Gene expression used to define cell types. C) UMAP of CD8 T cells colored by activation score (left) and exhaustion score (right). Scores were calculated using UCell with gene signatures for activation (NME1, IL2RA, CD38, HLA-DRA, HLA-DRB1, HLA-DRB5, MKI67, and PCNA) and exhaustion (CTLA4, HAVCR2, LAG3, PDCD1, TIGIT, and ENTPD1). D) Activation and exhaustion scores across cell types. E) UMAP of CD8 T cells from AML marrow (Desai et al, red, #cells=17,715) and solid tumor tumor infiltrating lymphocytes (Zheng et al, grey, #cells=184,455). F) UMAP of TOX, PDCD1, LAG3, and TIGIT gene expression in CD8 T cells. G) CD8A, PRF1, TOX, PDCD1, LAG3, and TIGIT expression in CD8 T cells across cancer types. BC=breast cancer, BCL=B cell lymphoma, CHOL=cholangiocarcinoma, ESCA=esophageal cancer, FTC=fallopian tube carcinoma, MM=multiple myeloma, OV=ovarian cancer, PACA=pancreatic cancer, RC=renal carcinoma, THCA=thyroid carcinoma, UCEC=uterine corpus endometrial carcinoma. H) Differential gene expression analysis between CD8 T cells from patients who achieved remission versus those who did not. I) Pathway enrichment analysis of upregulated genes from CD8 T cells from patients who achieved remission. Highlighted in red are pathways related to TNF response and T cell activation.

To contextualize AML-associated T cell states, including the rarity of traditionally defined dysfunctional phenotypes, we integrated publicly available datasets from a pan-cancer TIL atlas^29^ and single-cell AML BM resource^11^ (Figure 3E), building on prior cross-dataset comparisons^11^. Compared with CD8 TILs from solid tumors, AML-associated CD8 T cells exhibited markedly lower expression of exhaustion-related genes such as *TOX, PDCD1*, *LAG3*, and *TIGIT* (Figure 3F-G). These findings are consistent with previous studies and suggest that AML T cell alterations are distinct and could arise through mechanisms that differ from those described in solid tumors.

We then compared transcriptomic features of CD8 T cells from patients who achieved remission with those who did not. Differential gene expression analysis revealed that remission was associated with enrichment of pathways linked to TNF response and T cell activation. Key genes associated with remission included *NFKB1*, *NFKBIA*, *REL*, *RHOB*, *FOS*, and *GNLY* (Figure 3H-I). These results suggest that a preexisting, activated CD8 T cell program at diagnosis may support leukemic clearance.

### Viral-specific T cells cannot explain highly expanded T cell clones in AML patients

To investigate whether specific CD8 T cell phenotypes were associated with clonal expansion, we analyzed the TCR sequences from the 16 patient samples. Productive TCR alpha and beta chain sequences were recovered for 86% of CD8 T cells (n=20,275 cells), with comparable recovery rates across all clusters (range, 83.4-91.7%), indicating that TCR detection did not enrich for any CD8 T cell state (Supplemental Figure 5A-B).

We assessed the distribution of clonal expansion across phenotypes. As expected, the naïve CD8 T cell cluster exhibited the greatest TCR diversity, corresponding to lower clone frequency. In contrast, the most highly expanded clones were predominantly localized within the GZMK and GZMB TEMRA populations (Figure 4A-B). GZMK^+^ CD8 T cells have previously been reported to be enriched in high inflammation environments^43^. To quantify clonal overlap between phenotypic clusters, we calculated pairwise Jaccard similarity indices based on shared TCR clonotypes (Figure 4C). This revealed the greatest clonal overlap among effector memory, GZMK TEMRA, and GZMB TEMRA populations, while proliferating cells shared clones predominantly with the exhausted-like and GZMB TEMRA clusters. Overall clonal architecture varied substantially across patients, with five individuals harboring at least one highly expanded clone comprising more than 20% of their marrow CD8 T cell compartment (Figure 4D).

**Figure 4.**
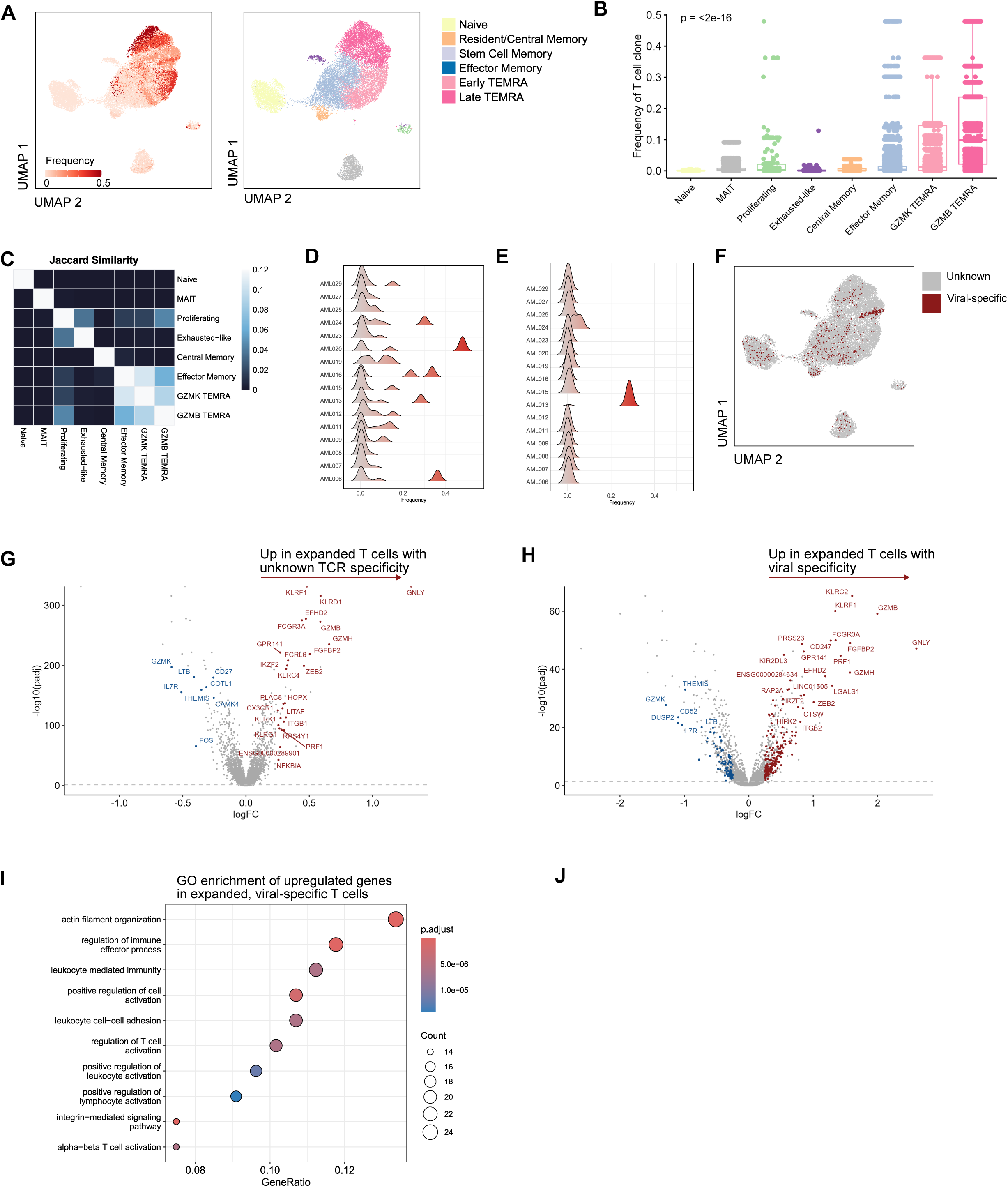
Viral-specific clones are insufficient to explain highly expanded clones in AML. A) UMAP of CD8 T cells colored by clonal frequency (N=16, #cells=20,275). For each patient, clonal frequency was calculated as the number of CD8 T cells assigned to a given TCR clone divided by the total number of CD8 T cells with a recovered TCR in that patient’s bone marrow sample. B) Frequency of clones across CD8 T cell types. Each point represents a T cell clone. C) Jaccard similarity index of CD8 T cell clones shared by cell type D) Clonal frequency distributions by patient. Right-shifted peaks reflect expanded clones. E) Clonal frequency distributions by patient, filtered on known viral-specific clones. F) UMAP of CD8 T cells colored by known TCR specificity. Red=viral-specific, grey=unknown specificity. G) Differential gene expression analysis between high clonal (>10% frequency) vs low clonal (<10% frequency and >2 cells) non-naive T cells with unknown TCR specificity. H) Differential gene expression analysis between high clonal (>10% frequency) vs low clonal (<10% frequency and >2 cells) non-naïve, viral-specific T cells. I) Pathway enrichment analysis of upregulated genes depicted in (H). J) Venn diagram of upregulated genes from (G) and (H).

Because healthy BM contains resident memory T cells, we next evaluated whether these expanded clones in AML could instead represent bystander viral-specific populations. To address this, we compared CDR3-TCRΒ sequences from our data to the VDJdb database^33^. We identified 622 CD8 T cells corresponding to 222 previously identified TCR epitopes (Supplemental Table 5). The most frequent matches were to epitopes derived from cytomegalovirus, Epstein-Barr virus, SARS-CoV-2, and influenza. Except for a single clone, all identified viral-associated clonotypes constituted <10% of the CD8 T cell repertoire within individual patients (Figure 4E). Additionally, viral-specific clones were distributed across multiple CD8 T cell clusters and were not enriched within the TEMRA populations (Figure 4F).

Given that viral-specific TCRs did not account for most expanded clones, we next investigated whether clonal expansion was associated with distinct transcriptional programs in viral-specific compared to T cells with unknown TCR specificity. We compared gene expression between highly expanded clones (>10% of patient CD8 T cells) and low-frequency clones (>2 cells, <10% frequency) within non-naïve, viral-specific and unknown TCR populations (Figure 4G-H, Supplemental Table 6). In both comparisons, highly expanded clones demonstrated upregulation of a shared cytotoxic effector program including expression of *PRF1, GNLY, GZMB, and GZMH*. However, viral-specific clonal expansion was associated with enrichment for genes involved in canonical TCR signaling and activation, including *ZAP70, CD247, TXK, VAV3,* and *FGR*, along with transcriptional regulators of effector differentiation such as *BATF* and *TBX21*. Gene ontology analysis of these genes confirmed enrichment of pathways related to T cell activation and actin cytoskeleton organization (Figure 4I). In contrast, clonally expanded T cells with unknown specificity were enriched for NK receptor genes (*KLRG1, KLRK1, KLRC4*), the transcriptional regulator *HOPX*, and the NF-κB pathway gene *NFKBIA* (Figure 4J). Together, we combined these five genes into a composite leukemia-associated NK-immune score (LaNKi), which we reasoned may capture this divergent, NK-like program of clonal expansion in AML. This NK-like, rather than canonical TCR-driven, program of clonal expansion prompted us to examine the utility of the LaNKi score more broadly in AML CD8 T cells.

### The LaNKi score identifies a distinct NK-like program enriched in AML CD8 T cells

To assess whether the LaNKi score could distinguish clonally expanded T cells of unknown specificity from other CD8 T cell populations, we examined three patients with highly expanded clonotypes. In patient AML006, the dominant clone comprised 36.2% of marrow CD8 T cells and lacked a match in the VDJdb database, indicating unknown specificity (Figure 5A). This clone was almost exclusively composed of GZMK TEMRA cells (Figure 5B) and exhibited a significantly higher LaNKi score compared to all other CD8 T cells in this patient (Figure 5C). A similar pattern was observed in patient AML020, where the dominant clone (47.9% of marrow CD8 T cells) was of unknown specificity, was predominantly GZMB TEMRA, and again showed a significantly elevated LaNKi score relative to background T cells (Figure 5D-F).

**Figure 5.**
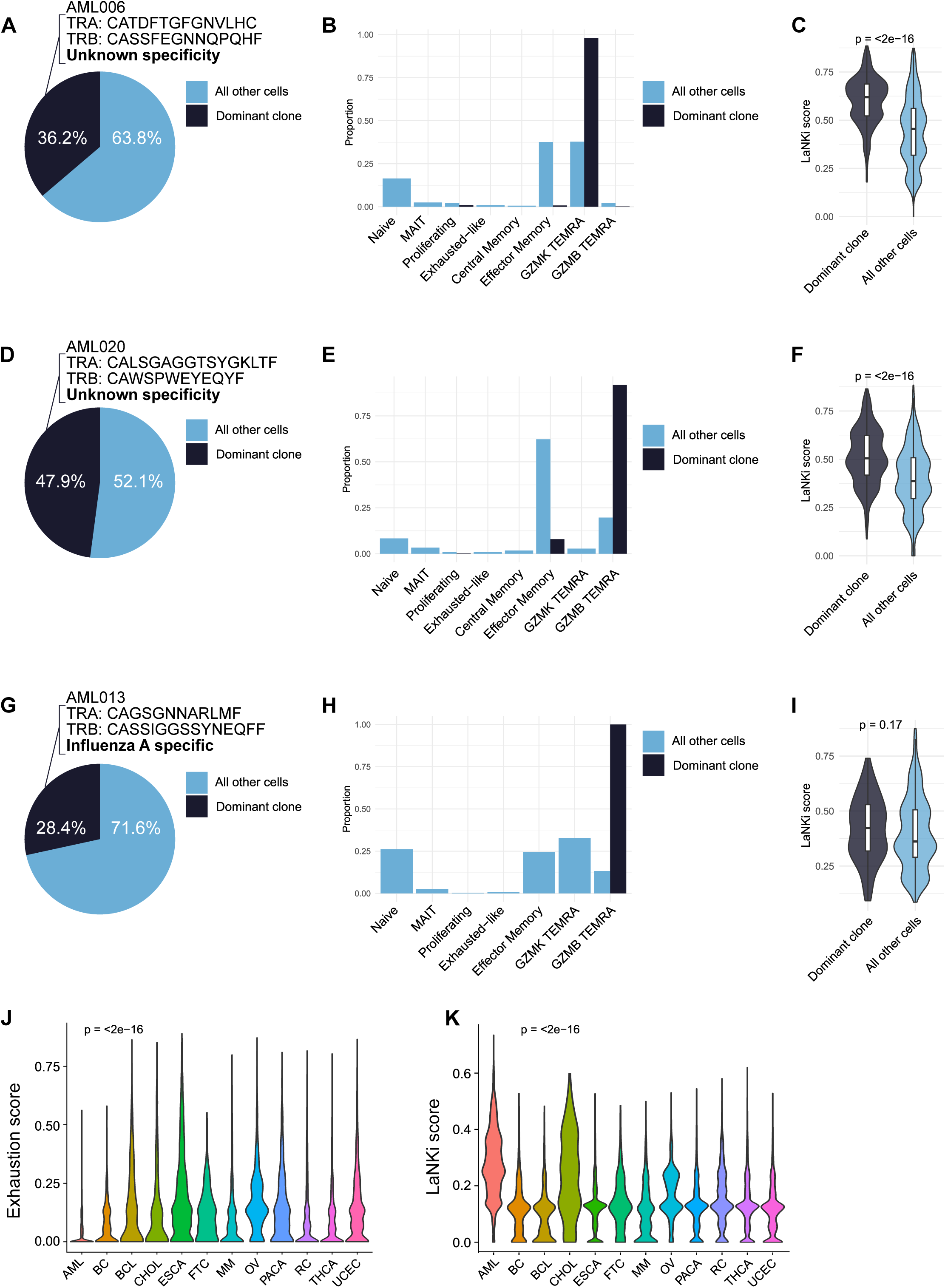
LaNKi score identified a leukemia-associated, NK-like CD8 T cell program distinct from viral-specific and solid tumor exhaustion states. A) Patient AML006: Pie chart of the clonal repertoire with the dominant clone (highest frequency) shown in dark blue and all remaining clones aggregated in light blue. This clone had unknown specificity. B) Patient AML006: Grouped bar plot showing the phenotypic distribution of the dominant clone (dark blue) versus aggregated background clones (light blue) across cell types. Bars for each clone group sum to 1 across all cell types. C) Patient AML006: Comparison of the LaNKi score in the dominant clone versus background clones. P-value calculated using Wilcoxon test. D) Patient AML020: Pie chart of the clonal repertoire with the dominant clone (highest frequency) shown in dark blue and all remaining clones aggregated in light blue. This clone had unknown specificity. E) Patient AML020: Grouped bar plot showing the phenotypic distribution of the dominant clone (dark blue) versus aggregated background clones (light blue) across cell types. Bars for each clone group sum to 1 across all cell types. F) Patient AML020: Comparison of the LaNKi score in the dominant clone versus background clones. P-vacalculated using Wilcoxon test. G) Patient AML013: Pie chart of the clonal repertoire with the dominant clone (highest frequency) shown in dark blue and all remaining clones aggregated in light blue. This clone was specific to Influenza A. H) Patient AML013: Grouped bar plot showing the phenotypic distribution of the dominant clone (dark blue) versus aggregated background clones (light blue) across cell types. Bars for each clone group sum to 1 across all cell types. I) Patient AML013: Comparison of the LaNKi score in the dominant clone versus background clones. P-value calculated using Wilcoxon test. J) Exhaustion score of CD8 T cells in AML and pan-cancer TILs (Zheng et al.). Exhaustion score was calculated in UCell using CTLA4, HAVCR2, LAG3, PDCD1, TIGIT, and ENTPD1. P-value calculated using Kruskal-wallis test. Post-hoc Dunn tests comparing AML to all other cancers resulted in p-values <0.05. K) LaNKi score of CD8 T cells in AML and pan-cancer TILs (Zheng et al.). LaNKi score was calculated in UCell using KLRG1, KLRK1, KLRC4, NFKBIA, and HOPX. P-value calculated using Kruskal-wallis test. Post-hoc Dunn tests comparing AML to all other cancers resulted in p-values <0.05. BC=breast cancer, BCL=B cell lymphoma, CHOL=cholangiocarcinoma, ESCA=esophageal cancer, FTC=fallopian tube carcinoma, MM=multiple myeloma, OV=ovarian cancer, PACA=pancreatic cancer, RC=renal carcinoma, THCA=thyroid carcinoma, UCEC=uterine corpus endometrial carcinoma.

To test whether the LaNKi score simply reflects the TEMRA differentiation state rather than clonal expansion of unknown specificity, we examined patient AML013, whose dominant clone (28.4% of marrow CD8 T cells) matched an influenza A-derived epitope in VDJdb. Despite comprising a similarly expanded, predominantly GZMB TEMRA population (Figure 5G-H), this viral-specific clone did not show a significant difference in LaNKi score compared to other CD8 T cells from the same patient (Figure 5I). This finding indicates that the LaNKi score is not a surrogate for TEMRA differentiation, but rather reflects a transcriptional program associated with clonal expansion of CD8 T cells with unknown specificity in AML.

We next asked whether this signature extended beyond our cohort. Using the pan-cancer TIL atlas^29^ and a single-cell AML BM resource^11^, we found that an exhaustion score calculated from canonical T cell exhaustion genes was low in AML CD8 T cells (Figure 5J), consistent with our earlier findings (Figure 3F-G). In contrast, the LaNKi score was highest among CD8 T cells in AML relative to solid tumor TILs, except for cholangiocarcinoma TILs which also had an elevated score (Figure 5K). Together, these data indicate that the LaNKi score captures a transcriptional program specific to clonally expanded T cells of unknown (and possibly non-viral) specificity in AML, distinct from both identified, viral-specific TEMRA expansion and canonical exhaustion in solid tumors.

## Discussion

We present a high-dimensional, multimodal analysis of CD8 T cell states in the BM of newly diagnosed AML patients, integrating mass cytometry, single-cell RNA sequencing, and TCR repertoire. Across 46 patients representing diverse AML subtypes, we observed a skewing of CD8 T cells toward terminally differentiated TEMRA states, with limited evidence of canonical exhaustion, expanding on prior results in a larger, independent cohort. Clonal expansion was most pronounced in TEMRA populations and known viral-specific bystander T cells could not account for these expansions. The enrichment of TEMRA cell states mirrors findings from studies of T cells in multiple myeloma, another marrow-based malignancy^44,45^. Subtype-specific AML phenotypes were associated with distinct CD8 T cell differentiation patterns. Genes related to response to TNF were upregulated in CD8 T cells from patients who achieved remission, and naïve T cell abundance correlated with overall survival. Collectively, these data help to define the AML marrow CD8 T cell landscape at diagnosis.

Our finding that AML-associated CD8 TEMRAs retain robust IFNγ and TNFα production in response to stimulation suggests that, despite their terminally differentiated phenotype and high CD57 expression, these cells are not intrinsically refractory to cytokine production or degranulation. Though, there was a considerable range among the AML samples. Some studies indicate that terminally differentiated CD8 T cells may exhibit reduced responsiveness to TCR engagement and greater reliance on natural killer-related pathways^41^. Indeed, we found that expanded clones of unknown specificity preferentially expressed NK-like receptors compared to viral-specific clones. These findings are consistent with a prior observation that AML-associated CD8 T cells diverge from an activation trajectory toward an NK-like phenotype^13^. Expression of NKG2D (*KLRK1*) on CD8 T cells has been shown to facilitate cytotoxic responses against tumor cells independently of classical peptide-MHC recognition^46^. This subset of expanded CD8 T cells in AML marrow may be activated through alternative recognition pathways, potentially including NK receptor-mediated responses to stress-associated ligands expressed within the leukemic niche. Specifically, NKG2D ligands MICA, MICB, and UL-16-binding proteins have been reported to be expressed on AML cells^47,48^.

To quantify this NK-like transcriptional program, we combined five genes enriched in clonally expanded, unknown-specificity T cells (*KLRG1, KLRK1, KLRC4, HOPX, and NFKBI*A) into a composite leukemia-associated NK-immune score (LaNKi). At the patient level, the LaNKi score was selectively elevated in dominant clonotypes of unknown specificity, but not in a dominant clonotype identified as influenza A-specific, despite comparable clone size and similar TEMRA phenotype. This finding suggests that the LaNKi score may reflect a transcriptional state specifically associated with clonal expansion of unknown, potentially leukemia-associated specificity, rather than terminal differentiation alone. When applied to CD8 T cells from an independent AML scRNA-seq cohort and a pan-cancer TIL atlas, the LaNKi score was among the highest in AML CD8 T cells relative to most solid tumor TIL populations.

Our study raises several important questions for future investigation. Although viral-specificity analysis excluded common bystander clones, we cannot directly confirm which of the highly expanded clones recognize leukemia-specific antigens or possess functional cytotoxicity. This question would be best answered by TCR antigen screening and epitope discovery^49,50^ but is beyond the scope of this study. Importantly, we cannot exclude the possibility that the unknown clones are still viral-specific. The LaNKi score may offer one path forward for prioritizing candidate leukemia-associated clones for such downstream antigen discovery efforts. The mechanisms driving TEMRA skewing and functional attenuation are also unclear, with potential contributions from AML-expressed ligands^51^, the AML cytokine milieu^52^, or leukemia-derived neoantigens^53,54^. Finally, the role of specific leukemia-associated antigens arising from translocations, inversions, or TP53 mutations in shaping T cell responses remains an open question with direct implications for immunotherapy design.

In conclusion, our study provides a comprehensive reference atlas of CD8 T cell differentiation, transcriptional programs, and clonal expansion in the AML BM at diagnosis. By integrating high-dimensional mass cytometry and single-cell transcriptomic analyses across a diverse cohort of AML patients, we report that the AML immune microenvironment is characterized by enrichment of terminally differentiated TEMRA populations and relative scarcity of canonically exhausted T cells. These findings challenge checkpoint-centric models derived from solid tumors. By linking leukemia phenotypes to T cell states and identifying immune features associated with remission, our work establishes a framework for stratifying AML patients by immune landscape. Finally, the transcriptional divergence we observe between viral-specific and potentially leukemia-specific clonal expansions, captured by the LaNKi score, suggests that CD8 T cell responses in AML may arise through distinct activation pathways involving both canonical TCR and NK receptor-associated signaling. Together, these findings establish a reference framework for understanding CD8 T cell biology in AML, provide a foundation for developing rational CD8 T cell-directed immunotherapies, and through the LaNKi score, highlight NK receptor-associated signaling as a candidate mechanism of CD8 T cell activation in the leukemia microenvironment.

## Data Sharing Statement

Computational analyses utilized previously published packages. Newly generated sequencing data will be available to reviewers at request and to the public at manuscript acceptance.

## Acknowledgements

The Vanderbilt University Medical Center VANTAGE Core provided technical assistance for sequencing, and VANTAGE is supported in part by Clinical and Translational Science Award Grant 5UL1 RR024975-03, Vanderbilt-Ingram Cancer Center Grant P30 CA68485, Vanderbilt Vision Center Grant P30 EY08126, and National Institutes of Health/National Center for Research Resources Grant G20 RR030956. YEC is supported by T32GM007347. RL is supported by the Mark Foundation for Cancer Research Endeavor Award, NIH/NCI (R01CA259480), American Cancer Society (RSG-22-036-01-DMC), and the Gabrielle’s Angel Foundation for Cancer Research. SCL is funded by NCI R01CA292932, Mark Foundation for Cancer Research Endeavor Award and Edward P. Evans Foundation Discovery Research Grant. RW is funded by the Mark Foundation Endeavor Award; R01HL150078 and 1PO1HL131477; and Edward P. Evans Foundation Discovery Research Grant. PBF is supported by a Mark Foundation for Cancer Research Endeavor Award, a Novartis Global Scholar Award, NIDDK R56 (R56DK138826), VA MERIT Award (I01BX005991), and a Gabrielle’s Angels Foundation for Cancer Research Award.

## Author Contributions

Conceptualization: YEC, PBF.; Methodology: YEC, PBF; Data collection: YEC, CRP, TG, CP, JAC; Data analysis: YEC, MAC; Resources: MRS, PBF; Funding acquisition: PBF; Project administration: PBF; Supervision: PBF; Writing: YEC, CRP, MRS, RL, SCL, RW, PBF.

---

**Supplemental Figure 1.**
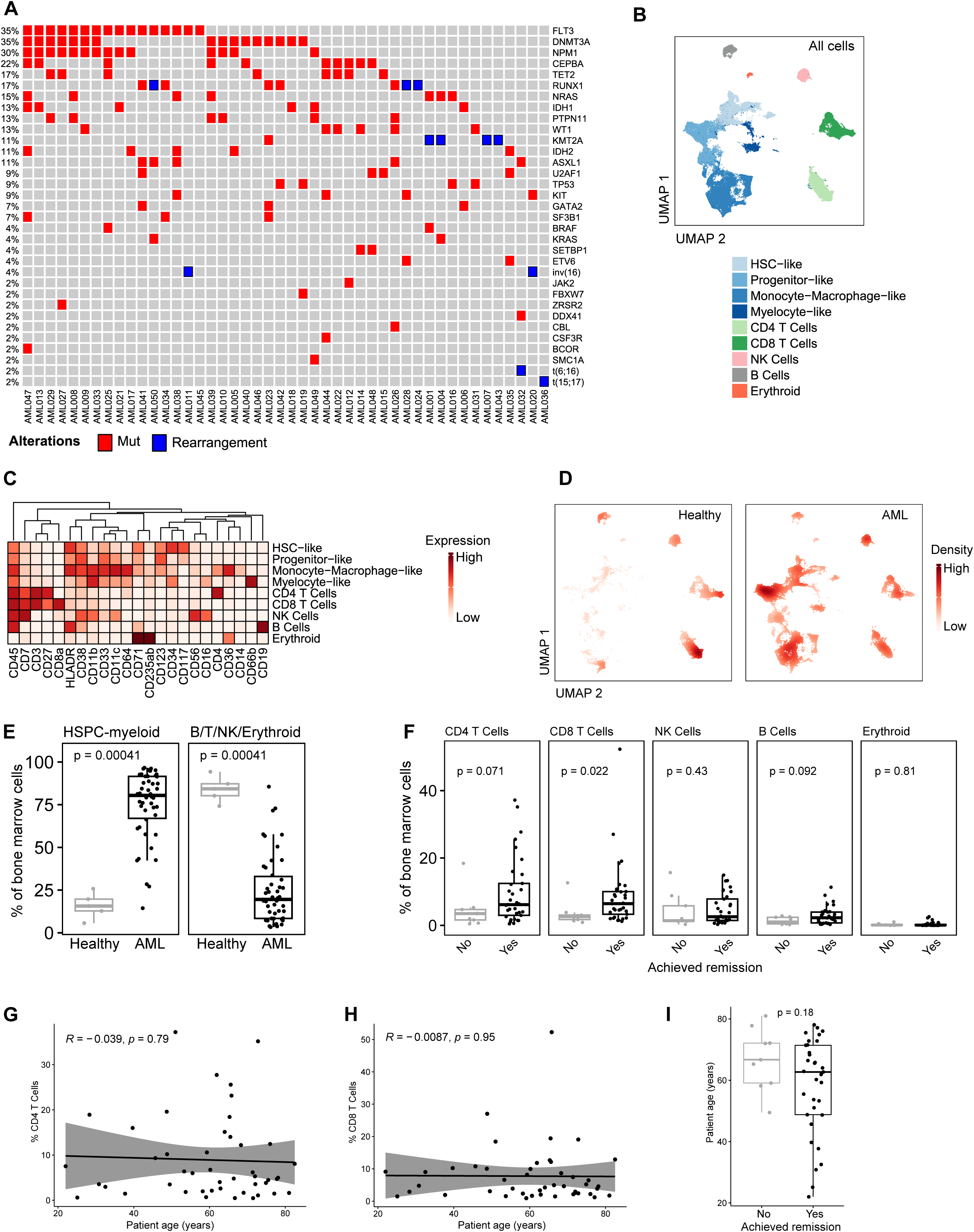
Immunophenotyping of the AML bone marrow microenvironment. A) Mutation status and cytogenetics of AML patient cohort. B) UMAP of bone marrow cells from all samples colored by cell type (N=51, #cells=658,917). C) Protein markers used to define cell types D) UMAP of cells from healthy (left, N=5, #cells=58,623) and all AML (right, N=46, #cells=58,623 randomly down sampled) marrows colored by density. E) Frequency of aggregated HSPC-myeloid (left) and B/T/NK/erythroid cells in healthy and AML samples. P-value calculated by Wilcoxon test. F) Frequency of cell types in AML samples from patients who achieved remission compared to those who did not. P-value calculated by Wilcoxon test. G) Pearson correlation between frequency of CD4 T cells and patient age in AML cohort. H) Pearson correlation between frequency of CD8 T cells and patient age in AML cohort. I) Comparison of patient age between those who achieved remission and those who did not. P-value calculated by Wilcoxon test.

**Supplemental Figure 2.**
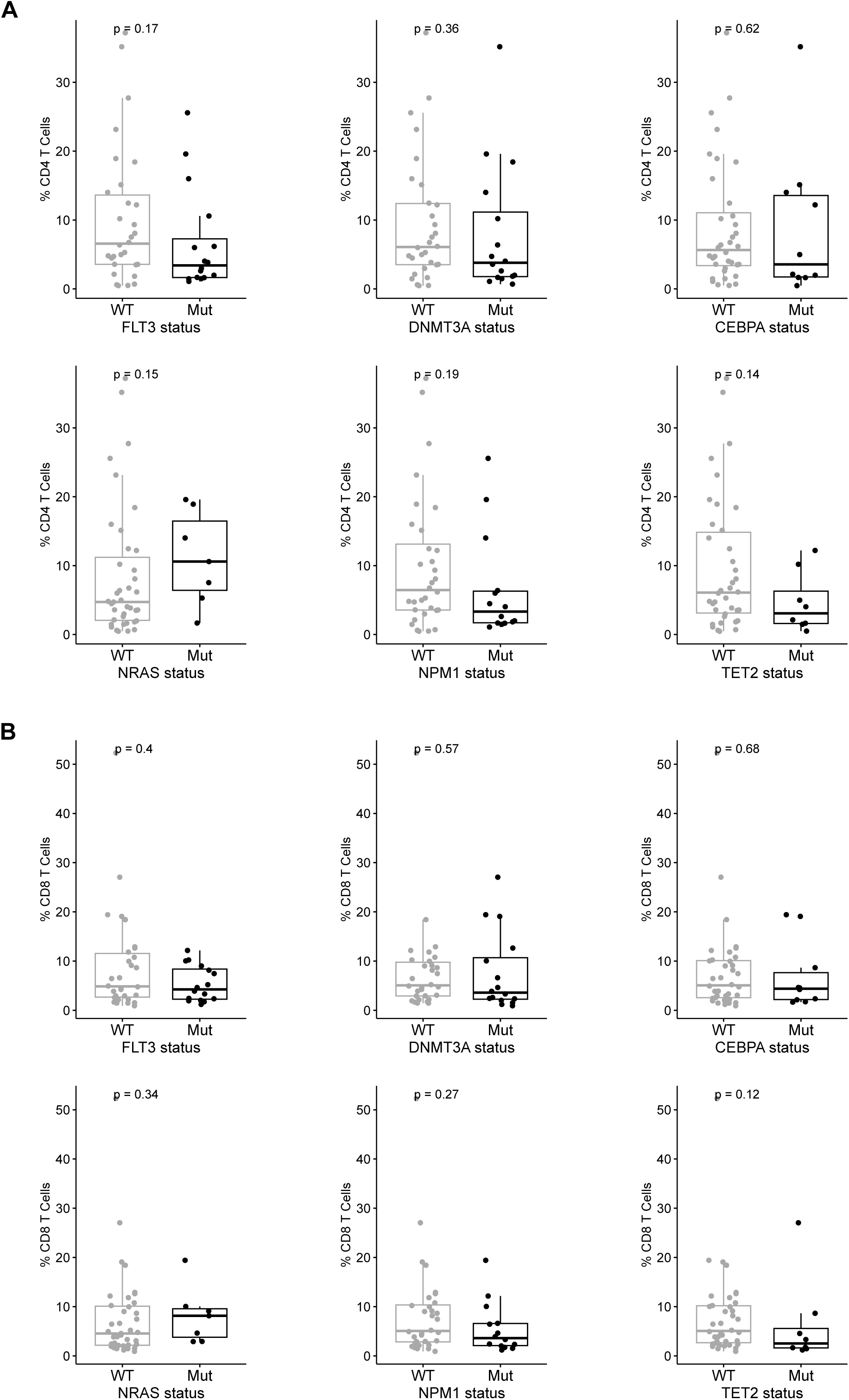
Mutation status is not associated with T cell abundance. A) Frequency of CD4 T cells in AML patients by FLT3, DNMT3A, CEBPA, NRAS, NPM1, and TET2 mutation status. P-value calculated by Wilcoxon test. B) Frequency of CD8 T cells in AML patients by FLT3, DNMT3A, CEBPA, NRAS, NPM1, and TET2 mutation status. P-value calculated by Wilcoxon test.

**Supplemental Figure 3.**
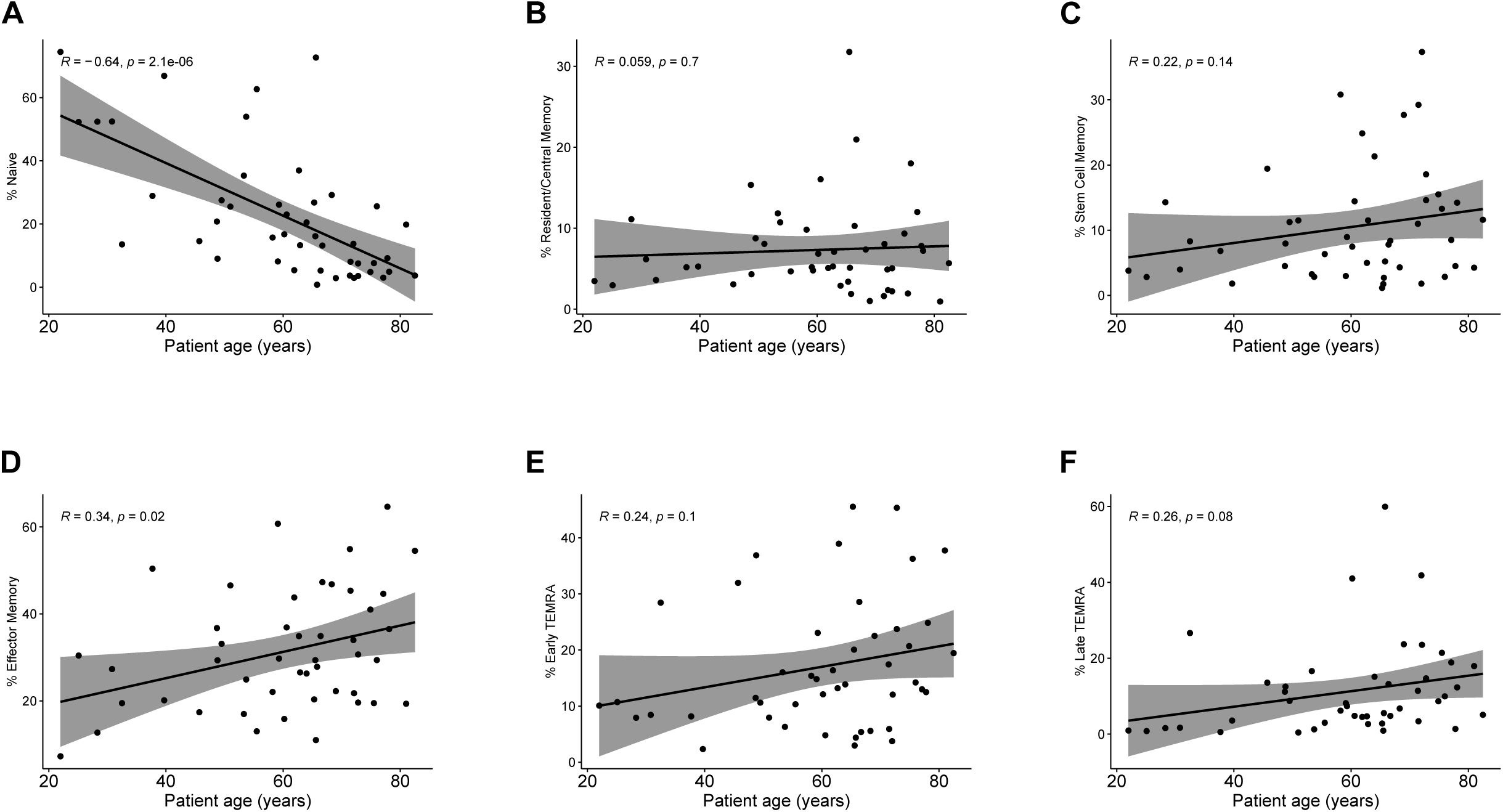
Correlation analysis between patient age and CD8 T cell phenotype. Pearson correlation between frequency of (A) Naive, (B) Resident/Central Memory, (C) Stem Cell Memory, (D) Effector Memory, (E) Early TEMRA, and (F) Late TEMRA and patient age.

**Supplemental Figure 4.**
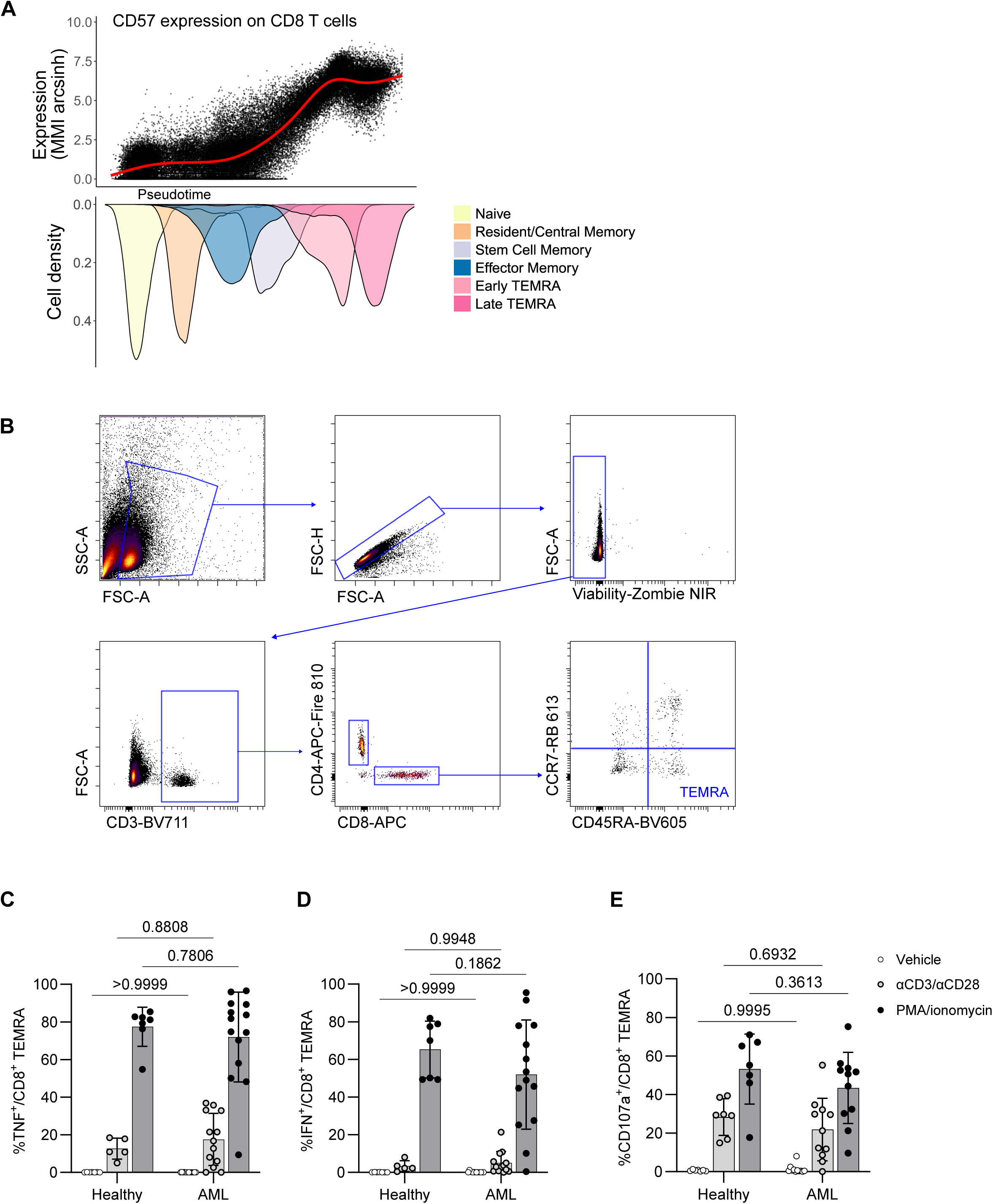
Functional testing of AML-associated CD8 TEMRA cells. A) Protein expression of CD57 in AML-associated CD8 T cells. MMI arcsinh = arcsinh transformed median mass intensity measured by mass cytometry. Mirrored density plots depict the dominant cell types across pseudotime. B) Gating scheme for CD8 TEMRA cells used for flow cytometry experiments. C) Percentage of TNFα positive CD8 TEMRA cells across healthy and AML samples stimulated with vehicle, αCD3/αCD28, and PMA/ionomycin. P-values calculated by Sidak’s multiple comparisons test. D) Percentage of IFNγ positive CD8 TEMRA cells across healthy and AML samples stimulated with vehicle, αCD3/αCD28, and PMA/ionomycin. P-values calculated by Sidak’s multiple comparisons test. E) Percentage of CD107a positive CD8 TEMRA cells across healthy and AML samples stimulated with vehicle, αCD3/αCD28, and PMA/ionomycin. P-values calculated by Sidak’s multiple comparisons test.

**Supplemental Figure 5.**
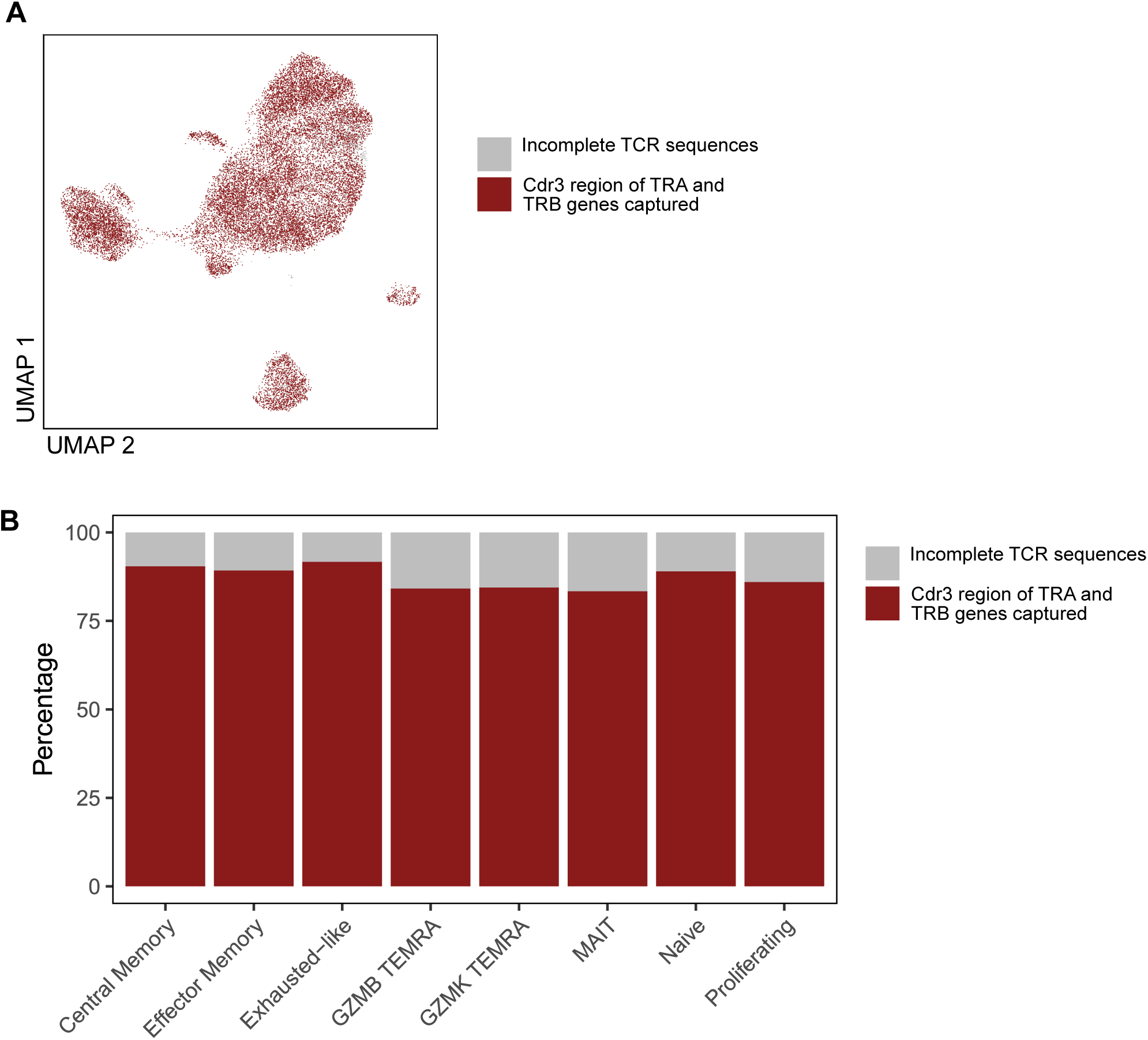
TCR sequence recovery across CD8 T cell clusters. A) UMAP of CD8 T cells colored by TRA and TRB cd3 sequence capture. B) Percentage of TRA and TRB cdr3 sequence capture across CD8 T cell types.

