## Supplemental Methods for "Clonally Expanded CD8 T Cells in AML Adopt an NK-Like Program Distinct from Canonical Exhaustion"

### Mass cytometry analysis

Flow cytometry standard (FCS) files were bead-normalized (CyTOF Software). Viable cells were manually gated in Cytobank (Beckman Coulter) based on cisplatin exclusion and iridium-DNA intercalator intensity. For the T cell-specific panel, viable CD8 T cells were further gated in Cytobank prior to export. Following gating, individual samples were debarcoded and compensated using the CATALYST package in R[19,20]. Signal intensities were harmonized across batches using CyCombine[21]. Dimensionality reduction was performed using Uniform Manifold Approximation and Projection (UMAP) on the corrected expression matrix.

Cells were clustered using the Louvain algorithm and annotated according to canonical lineage and differentiation markers. For trajectory analysis, pseudotime was inferred using the Slingshot package in R[22], designating the naïve CD8 T cell cluster as the root node.

### Single cell RNA and TCR sequencing

BMMCs were fluorescence activated cell sorted for viable, $CD45^{+}CD3^{+}$ cells. Individual patient samples were stained with TotalSeq-C antibody-derived hashtags (BioLegend) and pooled in equal proportions. Samples were stained with a panel of TotalSeq-C CITE-seq antibodies targeting cell surface proteins (Supplemental Table 2).

Pooled cells were loaded onto a 10x Genomics Chromium Controller using the Chromium Next GEM Single Cell 5' Reagent Kit. Reverse transcription, GEM cleanup, and cDNA amplification were performed according to the manufacturer's protocol. Amplified cDNA was split to generate parallel libraries for 5' gene expression, CITE-seq and hashtags, and TCR V(D)J enrichment. Libraries were sequenced on an Illumina NovaSeq X using 150bp paired end chemistry.

### Single cell RNAseq data processing

Reads from newly generated single cell RNAseq data were processed using Cellranger-v.9.0.1 (10X Genomics). For initial quality control of droplets, CellBender-v0.3.0[23] was performed. Then, cells with > 20% mitochondrial content and < 500 detected genes were filtered. Doublet detection was performed using scDblFinder[24] and cells identified as doublets were filtered. Count matrices were loaded into R-v4.5.1 and independent samples were concatenated into one Seurat-v5[25,26] object. Counts for transcripts were normalized using the *SCTransform*[27] function and counts for feature barcoding were normalized using the *NormalizeData* function with "*normalization.method* = *'CLR'*". Cell hashtags were demultiplexed using the *HTODemux* function with "*positive.quantile = 0.99*". For single cell immune receptor sequencing, contig annotations from CellRanger v9.0.1 were filtered for productive TCRs and imported into anndata objects as metadata. TCR frequencies were calculated as a proportion of the CD8 T cells from the respective patient.

**Dimensional reduction and data integration**

For integration of newly generated single cell RNAseq data, integration features were identified using the *SelectIntegrationFeatures* function with "*nfeatures=3000*". Integration features were scaled and PCA was performed using the *ScaleData* and *RunPCA* functions, respectively, with default settings. The *RunHarmony* function from Harmony-v1.2.3[28] was used to model batch effects with the sequencing lane as a categorical covariate. Nonlinear dimensional reduction was performed using Uniform Manifold Approximation and Projection (UMAP) with the *RunUMAP* function using *"spread = 1, min_dist = 0.5"*.

For integration of publicly available datasets, Zenodo Record 7883681[11] and GSE156728[29] were merged into a single Seurat v5 object. Then, the RNA layer was joined using the *JoinLayers* function. The *NormalizeData*, *FindVariableFeatures*, and *ScaleData* functions were used to preprocess the merged data. The *RunPCA* function was used for linear dimensional reduction and the *RunHarmony* function was used with the dataset as a categorical covariate. Finally, non-linear dimensional reduction was performed using the *RunUMAP* function with "*min.dist = 0.5*".

**Unsupervised clustering and cell annotation**

Unsupervised clustering was performed using the Louvain algorithm implemented in the *FindClusters* function with "*resolution=0.2*". The *FindAllMarkers* function was used to detect cluster-specific enriched genes. A consensus between cluster-specific gene markers and established celltype-specific markers were used to broadly annotate cells. Further subclustering of specific populations with various resolution settings ranging from 0.2 to 1 was performed to refine subcluster annotations.

**Differential expression and pathway enrichment analysis**

Differential expression was performed using the Wilcoxon rank sum test in Seurat-v5.1 with the *FindMarkers* function using the fast implementation by Presto[30]. Significance was determined by a log fold-change threshold > 0.25, an adjusted p value of < 0.05, and pct_diff > 5, where pct_diff = pct_in – pct_out, pct_in = the percent of observations in the group with non-zero feature value, and pct_out = the percent of observations out of the group with non-zero feature value. Pathway enrichment analysis was performed using clusterProfiler[31].
